# Fabrication & Characterization of Hyaluronic Acid/Eucalyptus Hydrogels Loaded with PLGA Nanoparticles with Methotrexate as an Injectable Therapy for Rheumatoid Arthritis

**DOI:** 10.1101/2025.02.18.638769

**Authors:** Maria Alejandra Castilla-Bolanos, Mariana Duenas, Laura Bustamante, Samuel Castillo

## Abstract

Rheumatoid arthritis is an autoimmune disease that affects about 250,000 Colombians, 82% of whom are women. Current treatments include disease-modifying antirheumatic drugs (DMARDs), such as methotrexate (MTX), analgesics and physiotherapy. The most common DMARD is MTX, which binds to plasma proteins with low efficiency (50%) and has a half-life of 6 hours. Due to its limitations when administered orally, nanoparticles (NPs) have been proposed to overcome these limitations. NPs support the release of therapeutic molecules, minimizing side effects and increasing the bioavailability of the drug in a controlled administration. NPs synthesized from biodegradable polymers, such as polyglycolic lactic acid (PLGA), are convenient for drug delivery due to their high biocompatibility and ability to bind DMARDs such as MTX. PLGA NPs loaded with MTX (MTX-PLGA-NPs) have reduced the presence of proinflammatory factors such as IL-10 and TGF-β, suggesting their potential as anti-inflammatory therapies for arthritis. Therefore, this study aims to develop MTX-PLGA-NPs in bioactive and biocompatible hyaluronic acid-eucalyptus (GelHA-E) hydrogels to preserve their stability and delay their degradation. PLGA-NPs were synthesized with an average hydrodynamic diameter of 200 nm, the 1237 cm-1 band found in FITR indicated the successful covalent conjugation with MTX; the mass loss of only 1% in GelHA-E indicated the thermogravimetric stability of the biomaterial and the low hemolytic and platelet aggregation percentage confirmed the biocompatibility of the biomaterial as a potential localized, anti-inflammatory, and injectable treatment for rheumatoid arthritis.

## I. Introduction

Rheumatoid arthritis is a progressive autoimmune disease characterized by the infiltration of inflammatory cells into the synovial tissue, which destroys cartilage and adjacent bone. The worldwide prevalence of rheumatoid arthritis has been estimated at 0.24% [1], and its prevalence in the United States and northern Europe may be even higher (between 0.5 and 1%) [1]. In Colombia, about 250,000 people suffer from arthritis and about 82% of these patients are women [2]. Although arthritis has no cure, there are palliative therapies that include disease-modifying antirheumatic drugs (DMARDs), such as methotrexate (MTX), sulfasalazine; and hydroxychloroquine [3]. In addition, there are biologics, such as tumor necrosis factor (TNF)-α inhibitors, interleukin-6 receptor blockers, and T-cell co-stimulatory blockers. Usually, this therapy is combined with analgesics, such as acetaminophen and opioids; and physiotherapy [4]. The most common of the DMARDs is MTX, which has a plasma protein binding efficiency of only 50% and has a half-life of approximately 6 hours [5]. Due to its short plasma half-life and low efficiency when administered orally, different types of nano vehicles have been proposed as alternative therapies to transport this drug more efficiently. The most common nano vehicles are liposomes and polymeric nanoparticles (NPs), which have been investigated for the transport and delivery of various drugs [6][7]. Nano vehicles are used to overcome drug efficiency limitations due to their ability to control the release of therapeutic molecules, reduce drug exposure to healthy tissues, minimize the side effects of drug overexposure, and increase drug bioavailability over time through a controlled delivery rate [6].

NPs can be made from different polymeric materials. For example, NPs can be synthesized from poly(lactic-co-glycolic acid) (PLGA), which is a biomaterial with great utility for drug delivery due to its high biocompatibility, biodegradability and ability to bind DMARDs such as MTX [7] [8]. Although MTX-loaded PLGA NPs (MTX-PLGA-NPs) have been mostly studied as an anticancer therapy, oncology studies have demonstrated the efficiency of PLGA-NPs to downregulate both proinflammatory factors and genes involved in the expression of these factors, such as IL-10, TGF-β, STAT3, and NF-κB, which would indicate their potential as an anti-inflammatory therapy for arthritis [8] [9]. Likewise, PLGA nanoparticles immobilizing MTX show properties suitable for specific targeted anti-rheumatic therapies [10], [11]. Furthermore, MTX-loaded NPs show adequate transfer to inflamed joints [10]. Another natural polysaccharide that has demonstrated significant potential as a drug carrier for various inflammatory diseases is Hyaluronic Acid (HA). It has been used in nano drug delivery systems for different diseases such as breast cancer, atherosclerosis, and rheumatoid arthritis (RA). This is primarily attributed to its high biocompatibility and biodegradability properties. Additionally, HA acts as a ligand for specific receptors present on the surface of the cells, such as CD44, which is implicated in RA tissue.

Moreover, HA has been physically and chemically integrated into various structures such as scaffolds, in-situ hydrogels, and viscoelastic solutions. Nonetheless, in many hydrogels, large pore sizes and high-water saturability have caused uncontrolled drug precipitation or rapid drug release. To overcome these issues, nanoparticles-based drug delivery systems have been introduced into hydrogels, leading to the formation of structures that offer multiple advantages, including increased drug loading capacity and sustained drug release.

Proinflammatory cytokines such as interlucin-6 (IL-6) and tumor necrosis factor-α (TNF-α) can cause injury to normal tissues by triggering inflammation anywhere in the human body. Excessive production of proinflammatory cytokines promotes many chronic inflammatory diseases, including rheumatoid arthritis. Extracts of Pistacia lentiscus [12] and Eucalyptus spp [13] are rich in phenolic compounds and show a strong reduction in IL-6 and TNF-α production, indicating strong anti-inflammatory effects, which is convenient to reduce the symptomatology caused by the disease.

Due to these therapeutic indications, in this study, MTX-PLGA-NPs were encapsulated in a hyaluronic acid hydrogel with - and without - eucalyptus (GelHA-E) to preserve the stability of MTX and to avoid its rapid degradation in a bioactive and biocompatible hydrogel as a vehicle. To evaluate the fabrication and use of NPs as nanocarriers, MTX-PLGA-NPs were synthesized using the double emulsion solvent evaporation method. Subsequently, the NPs were encapsulated in GelHA-E hydrogels to perform their chemical and biophysical characterization. PLGA NPs were synthesized with an average hydrodynamic diameter of 400 nm, and FTIR spectroscopy tests and thermogravimetric analysis (TGA) indicated MTX-PLGA chemical bonding. This research is a preliminary study describing the initial synthesis and characterization of PLGA NPs biomaterial and its loading with MTX for further pharmacodynamic evaluation and characterization of the anti-inflammatory effect of MTX-PLGA-NPs in GelHA-E for use as MTX-release therapy in the treatment of rheumatoid arthritis. In order to assess the effect of the inclusion of eucalyptus in the fabrication of an MTX-loaded hydrogel with PLGA nanoparticles, this study has the main objective to develop and characterize a gelatin-hyaluronic acid-eucalyptus gelatin hydrogel (GelHA-E) loaded with PLGA-methotrexate nanoparticles that enables the stability and release of methotrexate as a localized and effective treatment for rheumatoid arthritis.

## II. Materials and Methods

### A. Materials

Hyaluronic acid (HA) (53747-1G) and poly(lactic-co-glycolic) (430471-1G) were purchased from Sigma-Aldrich (St. Louis, MO, USA); Polyvinyl alcohol (PVA) (A2255) was purchased from Panreac (Barcelona, Catalonia, Spain); Methotrexate (2013M-0001140-R1) was purchased from TecnoFarma, S.A. (Bogota, D.C., Colombia); Gelatin (Gel) (FC-218567A) was purchased from Chem Cruz Biochemicals (Santa Cruz, CA, USA).

### B. Synthesis of PLGA-NPs y MTX loading

MTX-PLGA-NPs were prepared by the double-emulsion solvent evaporation method described by Maleki et. al.[14]. Briefly, MTX was dissolved in acetone (1mg/ml) and slowly added to the organic phase containing PLGA in chloroform (10 mg/ml). The obtained solution was sonicated for 1 minute (Branson M2800 ultrasonic bath, 20kHz). The polymer solution was added dropwise to an aqueous phase of PVA (0.1 % w/v) under stirring for 12 h (250 rpm). The NPs were separated by centrifugation at 6,500 rpm for 20 min and washed with type II water (3x). The suspension was flash-frozen at -80 °C for 12 h and freeze-dried under vacuum pressure below 50 m Torr for 48 h. Blank PLGA-NPs (without MTX) were prepared according to the same procedure. To identify the effect of PVA inclusion on size distribution and encapsulation efficiency, PVA parameters (0.05; 0.1; 0.2 % w/v) were investigated by a single factor experiment.

### C. Manufacture of GelHA & GelHA-E hydrogels

Gel (0.01% w/v) and HA (0.01% w/v) prepared in eucalyptus infusion (2 g/ mL) were combined in 4:1 ratio at 50°C under constant agitation for 30 min. Subsequently, MTX-PLGA-NPs (0.1 mg/mL) were added to the hydrogel.

### D. FTIR Spectroscopy

The surface modifications of MTX-PLGA-NPs and their conjugation to the biomaterial GelHA-E were analyzed by Fourier transform infrared spectroscopy (FTIR) (Bruker Optik GmbH, Ettlingen, Germany). The spectrum collected a range of 4,000-600 cm-1 with a spectral resolution of 2 cm^-1^.

### E. TGA thermogravimetric analysis

In order to know the stability of the biomaterial GelHA and GelHA-E with MTX-PLGA-NPs to temperature changes, a thermogravimetric analysis (TGA) was performed with 10 mg of samples (PLGA-NPs; MTX-PLGA-NPs; GelHA; and GelHA-E) in a TGA/simultaneous differential scanning calorimeter (TA Instruments, Newcastle, DE, USA). For the development of this characterization, samples were subjected to an increase in temperature at a rate of 10°C/min from 20°C to 500°C in a nitrogen atmosphere with a gas flow rate of 100 mL/min. The TGA data are shown as the percent mass loss as a function of temperature.

### F. Contact angle test

To measure the wettability and hydrophilicity of the samples, the contact angle test was carried out. For this, the drop-in-air or sessile drop method was used, due to its practicality compared to other methods such as the captive bubble method [15]. Since analytical and high precision and dosing instruments such as the OCA 35 [16] or OCA 15 EC [17] were not available, the assays were performed manually with a 1 mL syringe and a 26 × 76 mm slide. First, 2 mL of each sample was deposited in centrifuge tubes (Gel-HA; Gel-HA-E; MTX-PLGA-NPs-GelHA-E; and MTX-PLGA-NPs-GelHA). Using a 3 mL transfer pipette, 1 mL of the sample was deposited on the slide previously disinfected with alcohol. The syringe was then filled with type II water and positioned at a height of 5 cm above the slide, trying to obtain a 90° angle with respect to the slide. In this way, a drop of 2 μL was dropped and the moment when the drop fell on the sample was captured with the help of a SM-G975U camera (F1.5, 1/39s, 4.30mm, ISO640). This procedure was carried out at room temperature, trying to maintain the same height and angle in each attempt and performing 5 trials for each sample.

### G. Absorption/degradation test

Hydrogels are polymeric materials that absorb water into their structure without dissolving. Because of this, one of the most important properties is their swelling capacity. In order to know the swelling/degradation percentage of GelHA, GelHA-E, MTX-PLGA-NPs-GelHA, and MTX-PLGA-NPs-GelHA-E hydrogels, a preliminary absorption test was performed, which specified the amount of water that the hydrogel absorbed per sample. First, 0.6 mg of the freeze-dried samples of the three hydrogels were taken and immersed in buffered saline solution (PBS) at pH 7.4 and room temperature (23°C). The solution was filtered to retain the respective hydrogel and the hydrogel weight was measured every 30 min. Data collection was repeated for 120 minutes or until complete degradation of the hydrogel.

The swelling percentage was calculated according to equation 1:

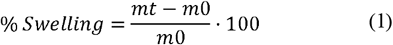

*mt* is the weight of the hydrogel at time and *m0* is the initial dry weight of the hydrogel. The equation is also used to calculate the maximum swelling of the hydrogel.

### H. Scanning Electron Microscopy (SEM)

Samples (MTX-NPs-GelHA and MTX-NPs-GelHA-E) were flash-frozen (−80°C) and freeze-dried (50 m Torr) for 48 h. This pre-preparation was carried out because the initial samples were liquid. Then, a coating with a thin film of gold particles was made on the surface of each sample to increase the electrical conductivity in them. Finally, they were taken to the SEM scanning electron microscope to evaluate their morphology and identify the e of the pores present.

### I. Sterilization and sterility testing methodology

In order to perform the biological characterization of the biomaterial, it is essential to sterilize it. For this purpose, the lyophilized samples (GelHA and GelHA-E) were subjected to ultraviolet (UV) exposure for 15 minutes per side, which allows sterilization of the surface of the material. Likewise, the liquid samples (MTX-PLGA-NPs-GelHA and MTX-PLGA-NPs-GelHA-E) were sterilized by filtration using a 0.22 µm filter.

Once sterilized, it is necessary to verify the absence of bacteria or microorganisms in the biomaterial. For this purpose, an analysis or sterility check by turbidity was used, consisting of the use of an appropriate medium for the proliferation of bacteria and microorganisms, using LB Agar medium. Subsequently, turbidity was observed qualitatively, and absorbance was measured after 24 h of incubation at 37° LB Agar was used as the experimental blank. [34]

### J. Antibacterial activity

GelHA and GelHA-E hydrogels were studied for their potential as an injectable vehicle for MTX-PLGA-NPs in joint tissue affected by chronic inflammation. Therefore, knowing their antibacterial properties is relevant to confirm their safe use in biomedical application. To know their antibacterial activity, agar diffusion methodology was carried out [32]. For this, LB Agar culture broth was prepared and polymerized in Petri dishes. Subsequently, reactivation of the bacterial strains, which were ultrafrozen (−80°C), was performed. The strains used were Escherichia coli (E. coli) and Staphylococcus aureus (S. aureus).

Subsequently, mass seeding of bacteria (E. coli and S. aureus; 0.001 absorbance) was performed in Petri dishes. The Petri dishes were divided into 5 areas for the evaluation of each type of hydrogel (MTX-PLGA-NPs-GelHA and MTX-PLGA-NPs-GelHA-E). In this way, the antimicrobial behavior of each type of biomaterial was evaluated. In the three missing areas, two antibiotics were added: one antibiotic that is highly effective against gram-positive microorganisms (Ampicillin) and another antibiotic that is highly effective against gram-negative microorganisms (Gentamicin).

### K. Hemolysis

According to ISO 10993-4, it is essential to evaluate the hemocompatibility of a material with tissues, especially with cells related to vascular pathways [18]. To carry out the hemocompatibility evaluation of the biomaterial, the study endorsed by ISO 10993-12 was performed, as presented in Table A1 (Appendix A) [19]. The hemolysis test helps to determine the capacity of a material to generate lysis in the main blood cells, the erythrocytes or red blood cells (RBCs). The procedure and hemolysis results of the GelHA and GelHA-E hydrogels evaluated in this test are shown below.

#### 1) Preparation of the extract

The necessary amount of PBS was calculated to obtain 5 mL of material extract (Table A1, Appendix A) [19]. The extracts were incubated for 24 h at 37°C with constant shaking (120 rpm). After this time, the extract was removed from the incubator and centrifuged for 15 min at 4,000 rpm. Because the material did not precipitate, it was necessary to filter the extract with a 0.22 µm syringe filter for use. In accordance with ISO 109993-12, it was determined that serial dilutions were necessary to obtain from an initial concentration (3 g/mL) a concentration endorsed by the standard (0.1g/mL) (Figure 2.8).

#### 2) Hemolysis protocol

At the same time, PBS buffer was prepared, and the erythrocytes were washed. For this, 4 mL of blood was added to an EDTA tube and then, the tube was centrifuged for 5 min at 1,800 rpm. Afterwards, the supernatant was discarded, and the same volume was again filled with NaCl solution (0.9% w/v). Then, it was mixed by immersion and taken to the centrifuge for 5 min at 1,800 rpm. This washing was repeated 3 times ending with the dissolution of the erythrocytes in PBS (10% v/v).

Finally, a 96-well microplate was used to carry out the controls of the experiment and to evaluate the hemolytic reaction of the material. Thus, a solution of the material extract (50 µL of Gel) was used as a positive control (P): as positive control (P) to a solution of material extract (50 µL of GelHA or GelHA-E) with erythrocytes (RBC) (50 µL) and triton (50 µL); as sample blank (BM) to a solution of PBS (50 µL) with material extract (50 µL of GelHA or GelHA-E); as negative control (N) to a solution of erythrocytes (50 µL of GR) with PBS (50 µL); and finally, as positive control blank (BP) to triton with PBS and negative control (BN) to PBS. Thus, to analyze the hemolytic effect of both extracts (GelHA or GelHA-E), 4 replicates (4 wells for each control) were performed to ensure the reproducibility and veracity of the experiment. Subsequently, the microplate was read in the spectrophotometer (450 nm) and the hemolytic activity of the material was evaluated.

### L. Data analysis

The data from each of the tests were processed in R v.4.2.1 software using the ggplot and plotly libraries. In addition, statistical comparison of the experimental groups was performed using ANOVAs and Kruskal Wallis according to the normal and nonparametric distribution of the data, respectively.

### M. Platelet aggregation

A platelet aggregation test was performed with the purpose of comparing GelHA and GelHA-E extracts with an aggregating substance (epinephrine). This test is necessary because when extracts of the hydrogels are obtained, they resemble the conditions in which the biomaterial will be in contact with the subdermal layers, so when they interact with the platelets it is possible to approximate this behavior with one that would be generated in a real physiological case. The principle of measuring platelet aggregation is based on the change in density between plasma with a high platelet count and plasma with a low platelet count when an agonist, in this case epinephrine, is added. Subsequently, when GelHA and GelHA-E hydrogels are added, the objective is to evaluate their potential to cause platelet aggregation. As there is a higher percentage of aggregation, the well becomes clearer and a greater amount of light is transmitted [20].

For the evaluation, 4 ml of blood was collected from a human donor in a tube with sodium citrate to prevent coagulation. Subsequently, it was centrifuged at 1,000 rpm for 10 minutes and the plasma was extracted. Then, using 2 mL tubes, the plasma was mixed in a 1:1 ratio with the sample extract (GelHA and GelHA-E), as well as with the positive control (epinephrine). Then, from each of these mixtures, 100 uL of sample per well was extracted into a 96-well microplate and read in the spectrophotometer at 450 nm. Finally, the potential of the sample as a platelet aggregator was identified. For this, the absorption values of the positive controls were compared with those of the samples.

### N. Zeta Size and Zeta Potential

PLGA-NPs hydrodynamic diameter and zeta potential were determined through DLS (Zeta-Sizer Nano-ZS, Malvern, UK) using five different pH (from pH of 4 to 8) to simulate the pH to which the biomaterial could be in contact in the body.

**Figure 2.**
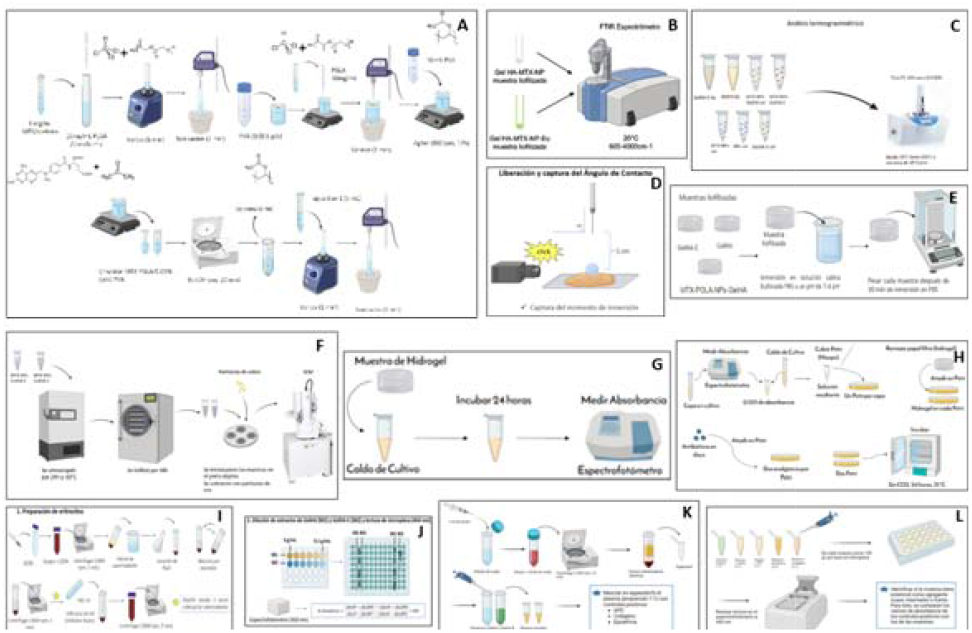
Materials and methods. **A**. Synthesis of MTX-PLGA-NPs. Procedure for synthesis of MTX-loaded PLGA nanoparticles. **B**. FTIR Spectroscopy. The FTIR spectroscopic test was performed in the spectrum 600-4000 cm^-1^. **C**. TGA analysis. Samples were placed in the TGA from 20°C a 500°C. **D**. Contact angle release and capture. 2 µL of distilled water was released over the sample and the contact angle was captured. **E**. Absorption test. The lyophilized samples were immersed in buffered saline solution at pH 7.4. The weight of each hydrogel was measured every 30 minutes (3). **F**. Scanning Electron Microscopy (SEM). The previous preparation of the (liquid) samples consisted of an ultrafreezing and freeze-drying process, as well as a gold coating to increase the conductivity of the samples **p**rior to SEM observation. **G**. Methodology for sterility testing. The sterility of the samples was observed qualitatively (turbidity) and quantified by measuring the absorbance of the samples after incubation in LB Agar medium (24 h, 37°C, without CO2). **H**. Antibacterial activity. The antimicrobial activity of GelHA and GelHA-E was evaluated in Petri dishe**s w**ith E. coli and S. aureus seeding. **I, J**. Hemolysis Methodology. [1] ISO, E. 10993-12: 2004. Biological Evaluation of Medical Devices—Part, 12, 10993-10912. **K, L. [**19] ISO, E. 10993-12: 2004. Biological Evaluation of Medical Devices—Part, 12, 10993-10912.

**Figure 3.1.**
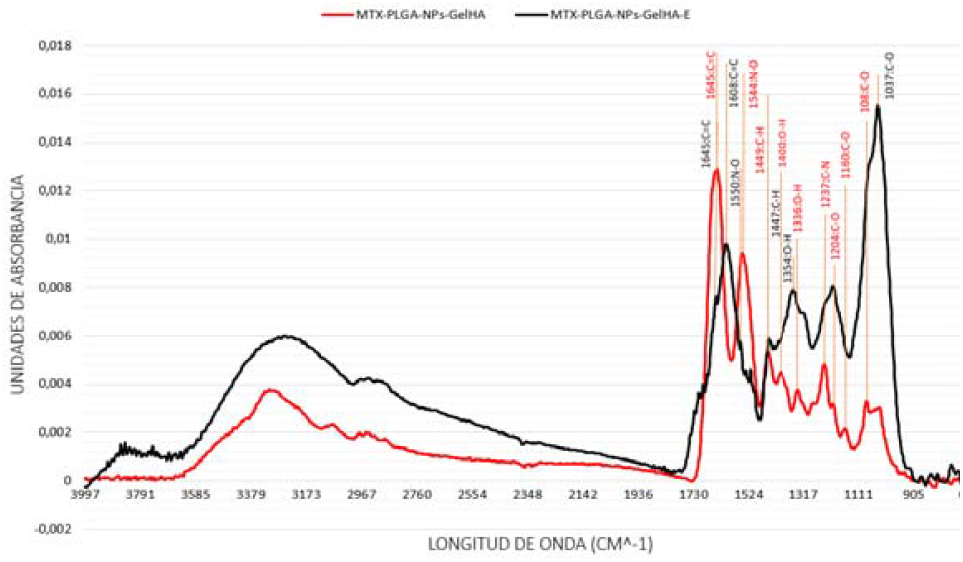
Absorbance vs. wavelength obtained from FTIR. Spectroscopic characterization identified the functional groups present in the MTX-PLGA-NPs-GelHA-E (black) and MTX-PLGA-NPs-GelHA (red) samples.

## III. Results

### A. FTIR Spectroscopy

Figure 3.1. shows the results obtained from the FTIR test for the MTX-PLGA-NPs-GelHA and MTX-PLGA-NPs-GelHA-E biomaterials. The points where the absorbance was highest and the respective associated functional groups (Figure 3.1) allow observing remarkable absorbance peaks at 1080, 1160, 1204, 1237, 1336, 1400, 1449, 1544, and 1639 cm^-1^. The other observable peaks do not present a marked relevance and cannot be concluded due to their low intensity or that they present less differences than the rest of the points in the graph. For the biomaterial MTX-PLGA-NPs-GelHA-E, which is distinguished by the inclusion of eucalyptus, the most notable absorbance peaks occur at 1037, 1204, 1354, 1447, 1550, 1608 y 1645 cm^-1^.

**Figure 3.2.**
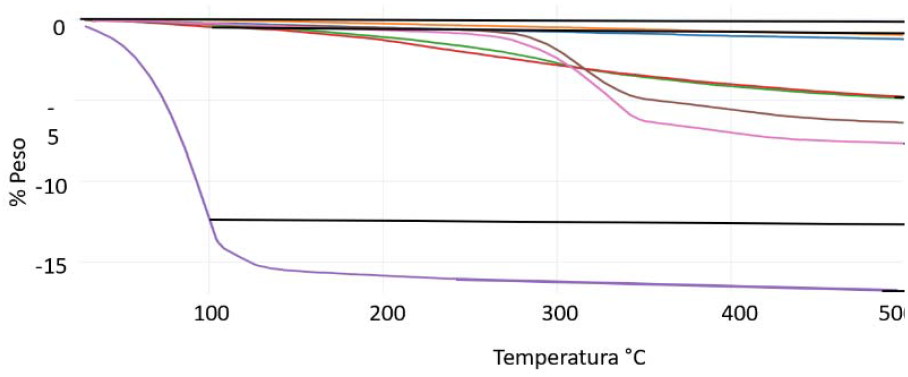
TGA thermogravimetric analysis. Weight % vs. temperature for GelHA, GelHA-E, MTX-NPs-GelHA-E, MTX-NPs, and NPs samples. Liq: liquid samples; Sol: solid (lyophilized) samples.

### B. TGA

Results obtained from TGA at 100°C show that freeze-dried MTX-PLGA-NPs-GelHA-E presents a more linear and stable curve than samples in the absence of eucalyptus, losing only 1% of its initial weight. In contrast, the freeze-dried MTX-PLGA-NPs-GEL-HA biomaterial decays more rapidly, losing approximately 13% by weight. In addition, it is evident that between both hydrogels in liquid state the results were very similar: at the end of the test both biomaterials lost only 1% of their weight, not highlighting significant differences (Wilcoxon test). On the other hand, the MTX-NPs and NPs samples presented a drop in weight near 230°C such that at the end of the test, at 500°C, MTX-NPs-sol decreased its weight by 6% and the NPs-sol sample by 8%.

### C. Contact angle

**Figure 3.3.**
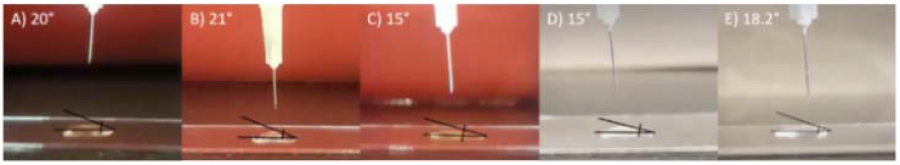
Average contact angle of GelHA hydrogels. Average contact angle of eucalyptus water: (A: 20°), GelHA: (B:21°), GelHA-E: (C: 15°), MTX-PLGA-NP-Gel-HA (D: 15°), and MTX-PLGA-NP-Gel-HA-E: (E: 18.2°).

### D. Absorption test

The initial sample weight of the hydrogels (GelHA-E, MTX-PLGA-NPs-GelHA, GelHA) was 0.6 mg. After performing the first cycle, the sample weights of GelHA, GelHA-E, and MTX-PLGA-NPs-GelHA increased to 2.5 mg, 1.8 mg, and 1.5 mg, respectively (Figure 3.1). The latter was dispersed in the PBS solution while GelHA and GelHA-E maintained their shape.

**Figure 3.4.**
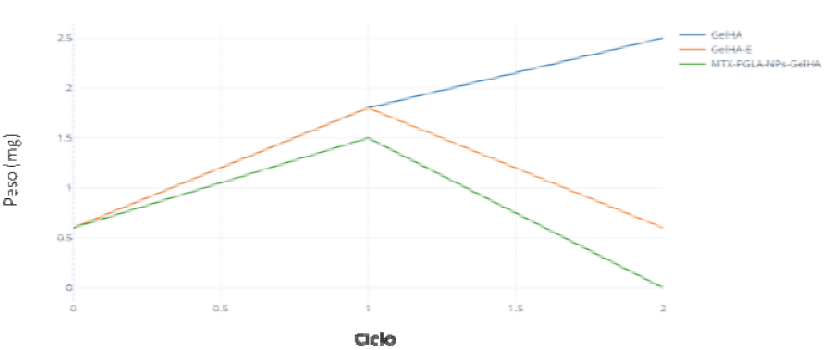
Weights of GelHA, GelHA-E, MTX-PLGA-NPs-GelHA hydrogels in cycles 1 and 2 of the degradation test. Blue: GelHA (mg); Orange: GelHA-E (mg); Green: MTX-PLGA-NPs-GelHA (mg).

At the end of the second cycle, the weight of the GelHA was 3.4 mg (Figure 3.2). The weight of the GelHA-E hydrogel was 0.6 mg, meaning that the hydrogel started to dissolve, a**s** it again dropped in weight (from 1.8 mg in cycle 1 to 0.6 mg in cycle 2). Finally, during the thirty minutes of the second cycle, th**e** MTX-PLGA-NPs-GelHA hydrogel was further dispersed, and at the end of the time it was completely dissolved.

The swelling percentage at the end of the first cycle was 316.6% for GelHA, 200% for GelHA-E and 150% for MTX-PLGA-NPs-GelHA. In contrast, once the second cycle was finished, the swelling percentage was 466.6% for GelHA and 0% for GelHA-E (because it returned to the initial weight). **B**ecause the MTX-PLGA-NPs-GelHA hydrogel dissolved, it was not possible to calculate its swelling percentage.

### E. SEM

In the case of the material that did not contain eucalyptus, a rather destroyed matrix can be observed with some structures that appear to be more preserved (Figure 3.5). These have a lamellar structure with pores up to 30 µm in diameter. In contrast to this, the material with eucalyptus instead of appearing destroyed presented itself as a compact, non-homogeneous and non-lamellar material (Figure 3.6). In addition, it did not present pores, although like the sample without eucalyptus, it did present a series of cavities. Also, in the sample with eucalyptus, the presence of small lighter clusters can be observed, so when studying the composition of these zones and comparing them with others where they were not present, it was found that these clusters contained percentages of Cl and the others did not.

**Figure 3.5.**
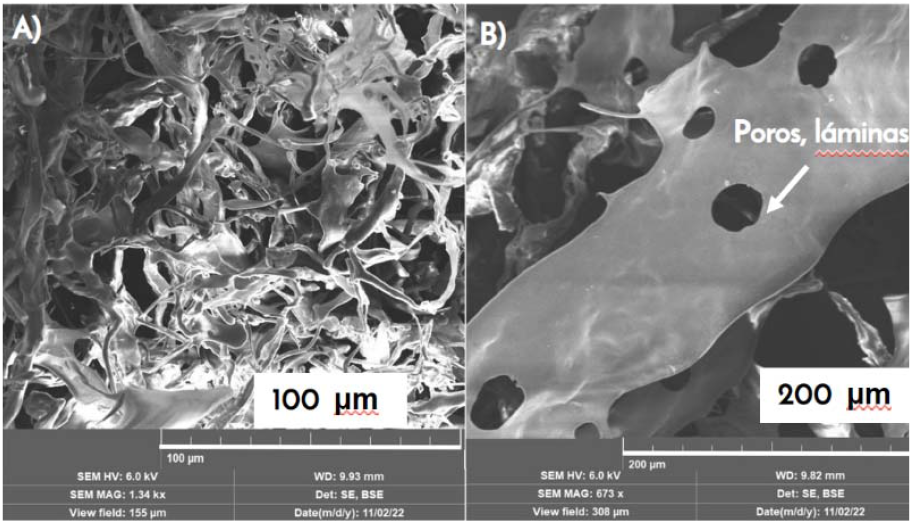
SEM of MTX-PLGA-NPs-GelHA. A) hydrogel matrix (100µm). B) preserved region. White arrow points to the presence of a pore.

**Figure 3.6.**
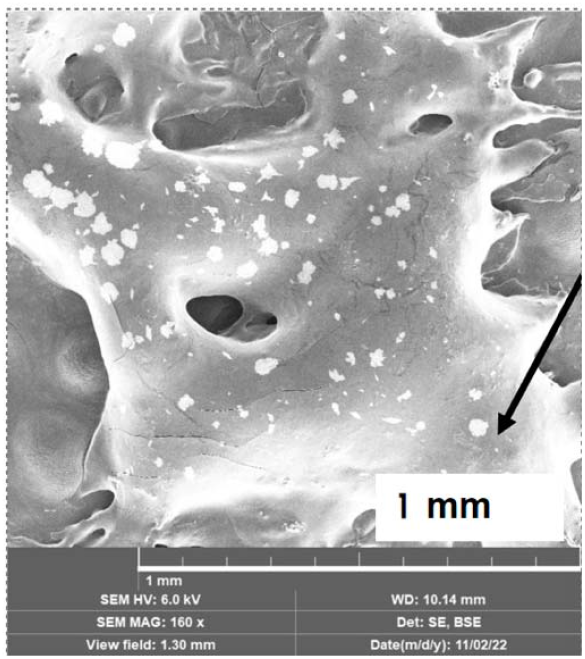
SEM of MTX-PLGA-NPs-GelHA-E. The black arrow shows aggregates composed of Cl.

### F. Sterility test

When observing the resulting absorbances, it can be stated that the absorbance between the LB Agar (negative control) and the samples does not show significant differences between them. And the overall absorbance values are low, close to the values of the empty samples (air only).

**Figure 3.7.**
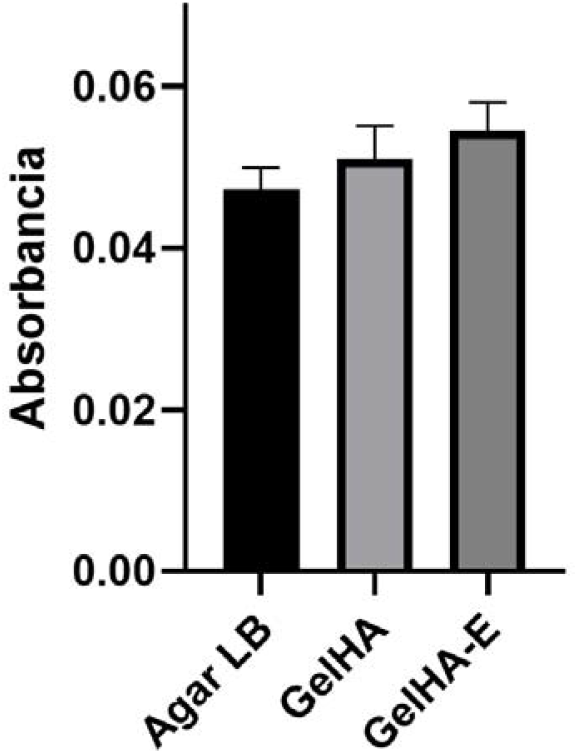
Absorbance of GelHA hydrogels.

### G. Antibacterial Activity

An inhibition halo around the antibiotics used (zones AB1 and AB2) shows that they fulfill their function, but the zones where the hydrogels are located (A and E) do not present any inhibition halo in any of the two Petri with strain.

**Figure 3.8.**
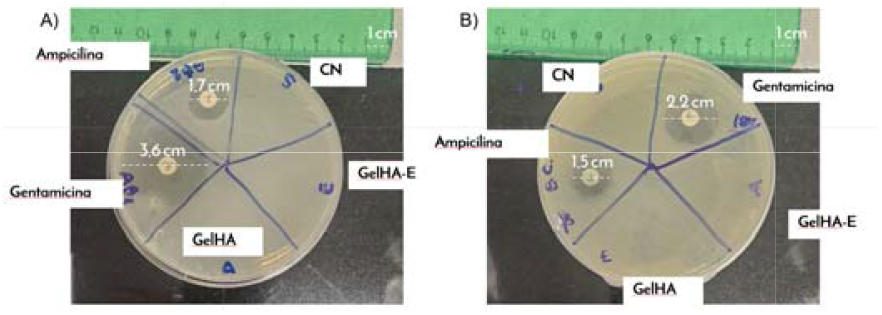
Antibacterial activity of *S. aureus* (A) y *E. coli*.

### H. Hemolysis

**Figure 3.9.**
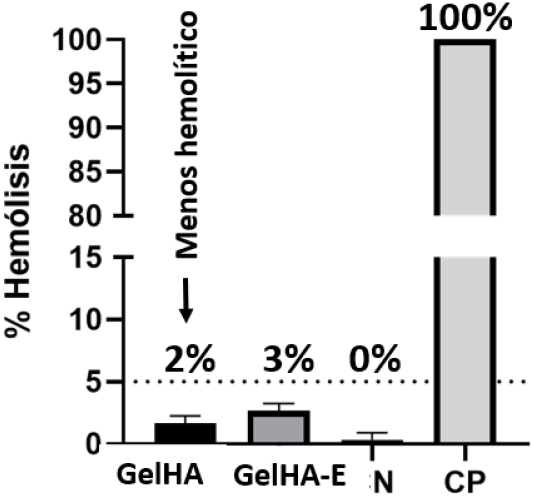
Hemolysis of GelHA (M1) y GelHA-E (M2); Negative control (CN), positive control (CP)

As evident from Table 2 and Fig. 3.9, all hydrogels (GelHA and GelHA-E) had a hemolytic percentage below 5%. The sample with the lowest hemolytic percentage was GelHA (2%). All percentages were compared with the positive control (100% hemolysis) which consisted of a solution of erythrocytes and triton.

**Table 1.** hemolysis of GelHA (M1) y GelHA-E (M2)

| M1 | M2 | CN | CP |
| --- | --- | --- | --- |
| 1% | 2% | 0% | 100% |
| 2% | 3% | 0% | 100% |
| 2% | 3% | 1% | 100% |

**Table 2.** Platelet aggregation of GelHA y GelHA-E.

| GelHA | GelHA-E | CN | CP |
| --- | --- | --- | --- |
| 6.00 | 6.02 | - | 100.00 |
| 6.60 | 6.36 | - | 100.00 |
| 7.02 | 6.45 | - | 100.00 |
| 7.79 | 6.53 | - | 100.00 |
| 6.85 | 6.34 | - | 100.00 |

### I. Platelet aggregation

**Figure 3.10.**
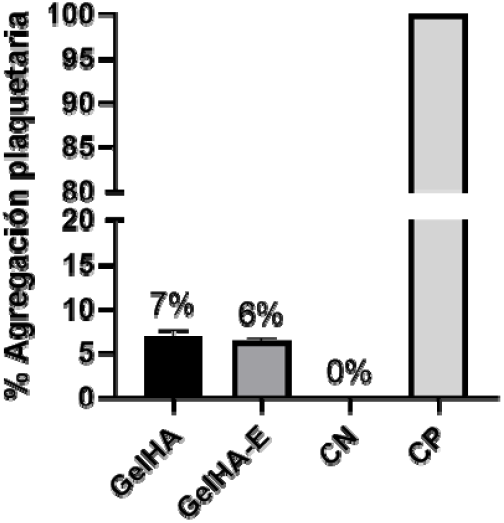
Platelet aggregation for GelHA y GelHA-E; negative control (CN); positive control (CP)

Table 3 and Figure 3.10 show the results of the percentage of platelet aggregation obtained from the GelHA and GelHA-E hydrogels. Accordingly, the two hydrogels presented a low percentage of platelet aggregation, less than 10% (6% GelHA-E, 7% GelHA). These percentages were compared with the positive epinephrine control (100% platelet aggregation).

**Table 3.** Zeta potential of GelHA and GelHA-E hydrogels.

| Sample | Diameter (nm) ± SD | PDI | Z-potential (mV) ± SD |
| --- | --- | --- | --- |
| PLGA-NPs (pH 4) | 231.667 ± 20.207 | | -4.030 ± 21.500 |
| PLGA-NPs (pH 5) | 295.000 ± 0.000 | | -2.670 ± 33.700 |
| PLGA-NPs (pH 6) | 281.667 ± 23.094 | | -1.610 ± 4.000 |
| PLGA-NPs (pH 7) | 255.000 ± 0.000 | | -1.910 ± 3.320 |
| PLGA-NPs (pH 8) | 268.333 ± 23.094 | | -2.210 ± 18.500 |

### J. Zeta Size and Zeta Potential

According to the results obtained from table 3, the sm size occurred at pH 4 with an average particle diameter of 231.667 nm. Conversely, the largest size was observe**d** at pH 5 with an average particle diameter of 295.000 nm. The lowest standard deviation (SD) in this measurement was foun**d a**t pH 5 and pH 7, indicating that at these pH levels, the particle size was more consistent and stable.

Additionally, all samples exhibited a negative Zeta-Potential, with the highest absolute value observed at pH 4 (4.03**0** mV), while the lowest was recorded at pH 6 (1.610 mV). In contrast to the Size results, the largest SD value was observed at pH 5 (_±_33.700 mV), whereas the lowest was seen at pH 7 (3.320 mV). This suggests that at pH 7, colloidal results appeared to be more stable.

**Figure 3.11.**
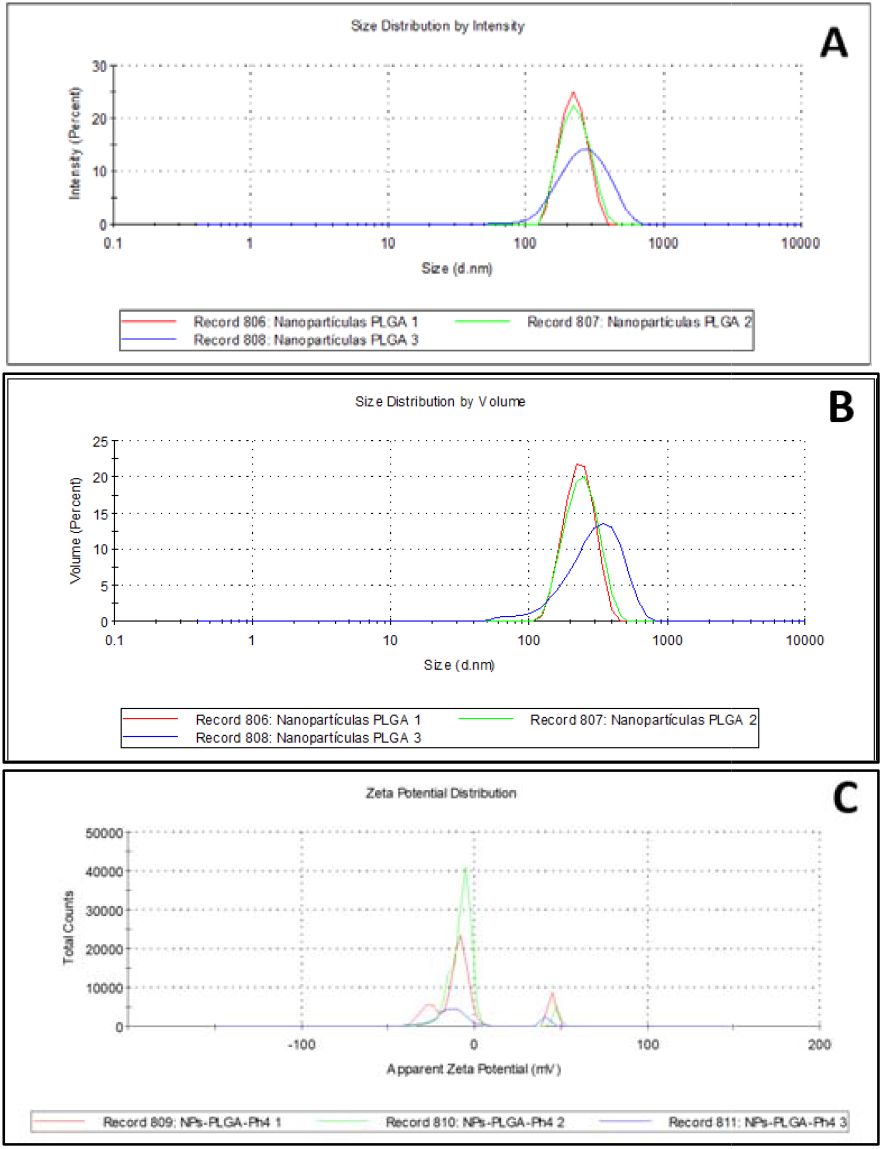
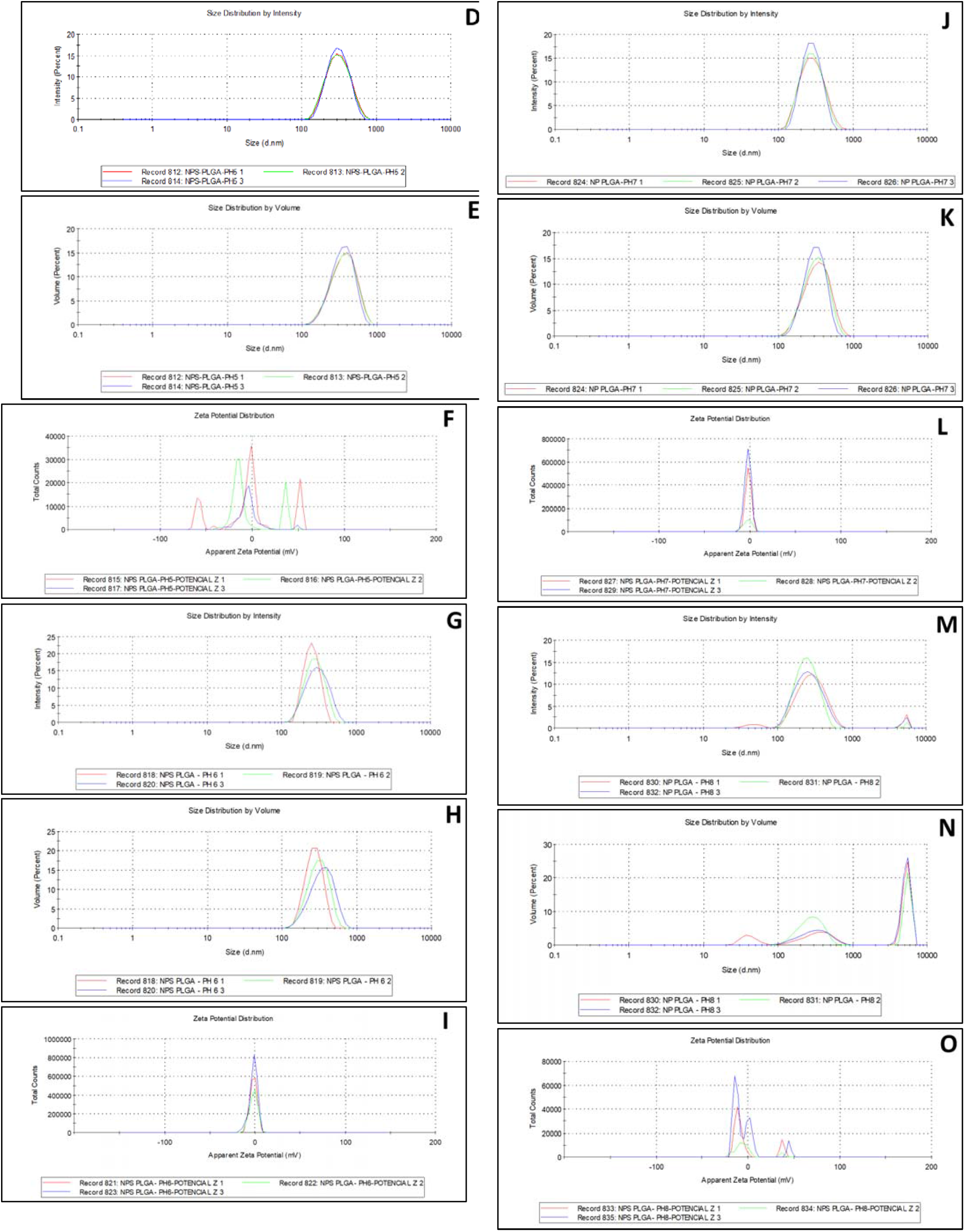
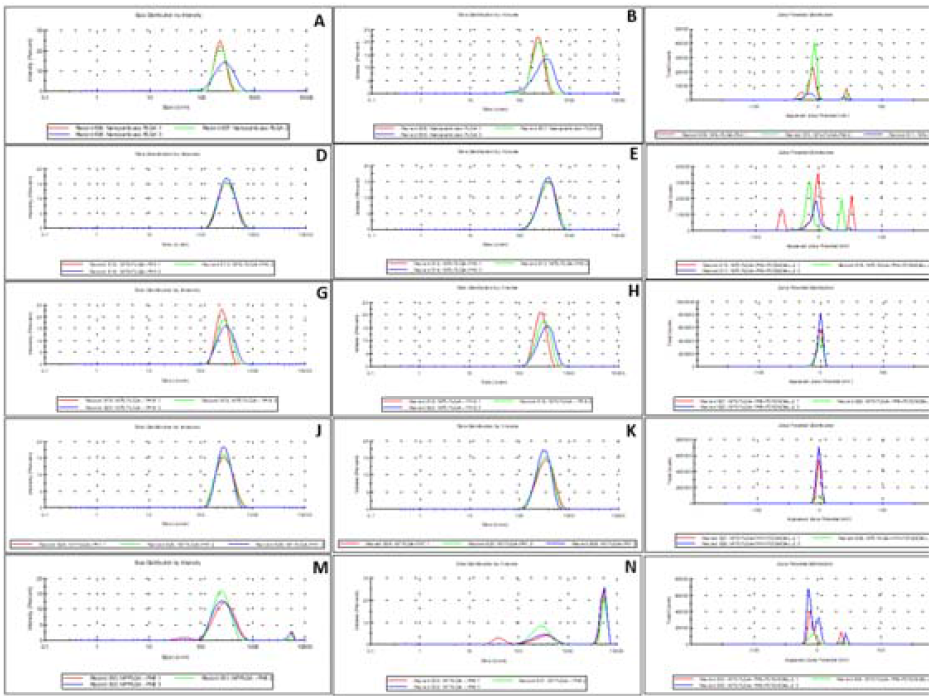
Zeta size and Zeta potential applying different pH values. **A, B, C. pH 4**. Size distribution by intensity (A), size distribution by volume (B), zeta potential distribution (C). **D, E, F. pH 5**. Size distribution by intensity (D), size distribution by volume (E), zeta potential distribution (F). **G, H, I. pH 6**. Size distribution by intensity (G), size distribution by volume (H), zeta potential distribution (I). **J, K, L. pH 7**. Size distribution by intensity (J), size distribution by volume (K), zeta potential distribution (L). **M, N, O. pH 8**. Size distribution by intensity (M), size distribution by volume (N), zeta potential distribution (O).

## IV. Discussion

This study developed GelHA hydrogels with MTX-PLGA-NPs and performed physicochemical characterization of the biomaterials based on their structure, thermal stability, hydrophilic and swelling behavior. The functional groups were obtained by FTIR spectroscopy. According to the results, the GelHA hydrogels possess a peak at1080 cm^-1^ hat is associated with C-O bonds of primary alcohols; a peak at 1160 cm^-1^ hat could belong to a C-O bond of tertiary alcohol; the peak at 1204 cm^-1^ is associated with the C-O bond of an ester, which is characteristic of HA. At 1237 cm^-1^, the bond at resonance is associated with an amine C-N bond, possibly from the drug MTX. The 1336 cm^-1^ resonance is associated with the O-H bond of PLGA, at 1400 cm^-1^ t can also be associated with an O-H bond but related to a carboxylic acid. Also, 1449 cm^-1^ may be due to the presence of C-H bonds with methyl groups, since it is found near 1450 cm^-1^. In addition, 1544 cm^-1^ can be related to the N-O bond and 1639 cm^-1^ to a C=O bond, or a C=C bond. [21]. Then, as the biomaterial without eucalyptus presents functional groups of HA, MTX and PLGA. it can be inferred that there was a correct reaction of the materials. As for GelHA-E, it seems to have many similarities in the absorbance peaks, although not necessarily in the exact wavelength: the peak at 1037 cm^-1^ may be the least accurate association: a primary alcohol C-O bond or even a C=C double bond is observed, at 1204 cm^-1^ an ester C-O bond, at 1354 cm^-1^ a O-H bond, at 1447 cm-^1^ it can also be approximated to a C-H methyl group, at 1550 cm^-1^ an N-O bond, and 1639 cm^-1^ a possible C=C bond. There is a notable difference at 1608 cm^-1^, which can also be associated with a C=C at 1639 cm^-1^[1].

Comparison of both materials reveals subtle s ectral differences, although they do not exhibit identical absorbance peaks. This may be attributed to peak coalescence, making individual peaks difficult to discern. For instance, near 1000 cm□^1^ in GelHA-E, the spectrum appears to merge with the peak at 1080 cm□^1^ observed in GelHA. Despite these minor variations, GelHA-E contains the chemical groups characteristic of the hydrogel components and MTX-NPs, suggesting that both materials correspond to the expected composition and confirming the success of the reaction process. The most relevan**t** bonds supporting this conclusion are the C-O bonds of alcohols in PLGA and hyaluronic acid, the O-H bond of hyaluronic acid, and the C-N amine bond of MTX. Furthermore, contact angle tests (Figure 3) confirmed the hydrophilic nature of the samples. Based on the test results, all five samples exhibited hydrophilic behavior [22]. with average contact angles ranging between 15° and 21°. The models demonstrating the highest hydrophilicity were Gel-HA-E and MTX-PLGA-NPs-GelHA, both with an average contact angle of 15°. Conversely, GelHA showed the lowest hydrophilicity with an average angle of 21°, followed by Eucalyptus extract (E) at 20°.

The results suggest that eucalyptus does not directly determine the hydrophilicity of MTX-PLGA-NPs-GelHA-E samples. However, the conjugation of GelHA with MTX-PLGA-NPs and MTX-PLGA-NPs-E exhibited higher hydrophilicity compared to GelHA and Eucalyptus extract alone. This effect aligns with previous studies on GelHA, where the contact angle decreased when combined with other polymers such as PDMS or PLGA. This phenomenon may be attributed to increased surface roughness resulting from crosslinking, which can lead to enhanced hydrophilicity.

On the other hand, a study by Souza et al [23] on the processing of gum arabic-based hydrogels indicates that, the addition of compounds such as eucalyptus and pine decreased the degree of swelling and absorption of the polymers because they caused a higher degree of crosslinking of the hydrogel [23]. In general, several studies show the great antioxidant, antibacterial, and anti-inflammatory properties of eucalyptus [24], as **we**ll as its penetration properties of bioactive molecules [25], but few show its effects at the level of hydrophilicity and how thi**s** changes according to the degree of hydrogel crosslinking. Likewise, it is considered that the range of error in the experiment was high because there were no high precision or dosing instruments such as the OCA 35 [26] or the OCA 15 EC [17] nor high resolution cameras that clearly and accurately captured the moment when the water droplet touched the surface.

Additionally, the TGA test yielded precise outcomes concerning the decomposition of both MTX-NPs and NPs samples. Previous research has indicated that MTX undergoes decomposition at 230°C, while PLGA experiences sig**n**ificant weight loss within the temperature range of 212°C to 600°C. Significant differences between freeze-dried MTX-NPs-GelHA-E and freeze-dried MTX-NPs-GelHA. Eucalyptus emerged as a key factor influencing the thermal stability of the sample. MTX-NPs-GelHA-E exhibited notably higher thermal stability compared to the variant without eucalyptus. The material lacking eucalyptus experienced rapid degradation, losing 0% to 13% of its weight between 20°C and 100°C. In contrast, the MTX-NPs-GelHA-E only lost 1% of its weight under the same conditions. This outcome and the one obtained in liquid samples (Fig. 3.4) provide compelling evidence of the enhanced thermal stability conferred by HA and Eucalyptus to the MTX-PLGA-NP structure. These favorable results align with previous studies showcasing the effective functionalization and stabilization of NPs composed of chitosan or curcumin/quercetin through HA encapsulation. Also, these outcomes can be attributed to the aforementioned study [23], which demonstrated that eucalyptus contributed to a higher degree of crosslinking within the hydrogel, imparting superior resilience to the material.

Nonetheless, thanks to TGA it can be determined that NPs were successfully encapsulated in HA, as. This is confirmed by other studies where HA improved the thermal stability of NPs.

Then, from the TGA results it is inferred that it would be better to use the biomaterial containing eucalyptus because in addition to the beneficial properties it provides, at room temperature and skin temperature it will be in a temperature range in which it will be stable and will be a better option for drug release since it will not have begun to degrade before application. Furthermore, it can be said that, while in all samples, near there 100°C there is a notable reduction in weight percentage due to water and moisture loss, in the case of MTX-NPs-GelHA-sol this weight percentage is higher [27]. The problem results in that the hydrogel are mostly composed of water so that at the time of application it may have lost many of its gel properties and may no longer be suitable for transdermal application due to a new consistency or also, that at the time of application being so degraded it cannot release the drug correctly and is released too quickly [28].

Based on the results of the preliminary absorption test, it was observed that the samples of the three gels presented different swelling percentages. GelHA was the one that obtained the most adequate results since as the cycles progressed its absorption percentage increased more than the MTX-PLGA-NPs-GelHA and GelHA-E gels, reaching 466.6%. This means that GelHA is the most resistant hydrogel to degradation. As for GelHA-E hydrogels, it showed a high degradation rate, since at the end of the second thirty minutes of the absorption test it returned to its initial weight of 0.6 mg. According to these results we can interpret that, although GelHA-E provides anti-inflammatory, antioxidant and antibacterial properties to drug delivery, to be used it requires its crosslinking with chemical agents, such as EDC to prevent its degradation in aqueous media. Finally, the MTX-PLGA-NPs-GelHA hydrogel proved to be the hydrogel with the highest degradation rate. This result was obtained during the first cycle, in which the erosion in volume of the hydrogel was observed, as well as its complete solubilization in the PBS solution in 1 h. However, the results obtained with the MTX-PLGA-NPs-GelHA hydrogel were not very different from those obtained with the GelHA-E hydrogel, suggesting that the best non-crosslinking option, according to its swelling, absorption and degradation behavior, for use as a drug delivery vehicle, is the GelHA hydrogel. This is arguably due to the fact that the pharmaceutical properties of hydrogels loaded with aqueous eucalyptus leaf extract may exhibit shear thinning behavior [29].

Moreover, the covalent integration of hydrogel particles in a secondary HA network produces structured and mechanically resistant hybrid networks [17], if hyaluronic acid is connected to other macromolecules or substances with lower biocompatibility, such as methotrexate or eucalyptus, characteristics would be lost, producing a hydrogel less resistant to degradation. Now, it is considered that the experimental procedure has a high range of error, since the instruments used, such as analytical balances, do not have adequate precision for measuring samples in milligrams (Adventurer Analytical OHAUS Model AX 124/E Balance, USA, error: 0.1 mg). Therefore, in future experiments, initial samples of greater weight will be taken in order to have more precise measurements of the absorption tests. Also, chemical crosslinking of the hydrogel will be performed to increase its resistance to degradation and provide a hydrogel with better swelling and degradation performance to be used as a drug delivery vehicle for arthritis.

In the case of SEM, taking into account that the samples had to be frozen and freeze-dried to perform the characterization, it was expected that the hydrogel matrix would degrade since the vacuum was extracted from the samples during freeze-drying and therefore the matrix fibers were destroyed. However, from some fragments that were not destroyed, it can be said that the initial morphology points to a lamellar structure. As for the pores, it is not possible to define with certainty whether these were present before the samples were frozen and freeze-dried because, in fact, in a study carried out on the morphological changes of hydrogels before and after the freeze-drying process it is mentioned that, although the initial hydrogel presented pores, when it was freeze-dried more pores were generated and of different sizes [30]. On the other hand, in the case of eucalyptus, a more condensed matrix without pores was found, which corresponds to that found in the literature, given that other studies have shown that adding eucalyptus to hydrogels altered their structure and prevented the formation of pores [20]. Also, the presence or absence of pores provides information on the mechanical strength of the material, so in this case the biomaterial with eucalyptus would have greater strength than the one without. This is due to the fact that the mechanical resistance in hydrogels increases as the size and presence of pores decrease [20]. This is appropriate because mechanical resistance is necessary in controlled drug release processes such as in this type of applications [20]. Likewise, the presence of clusters mentioned above given its composition with Cl must correspond to cross-contamination with surfaces impregnated with Cl atoms probably due to improper handling of the samples. This is because none of the materials used contained Cl.

According to the zeta potential and zeta size results obtained in Figure 3.11, it is evident that the size of NPs was not directly related to pH changes, as size values varied with the increment of pH, indicating that pH was not a direct determinant of PLGA-NPs size. The smallest diameter (231.667 nm) was observed at pH 4; however, it had a high standard deviation (±20.207 nm), implying greater variability in the size of particles. On the other hand, the second smallest size was seen at pH 7 (255.000 nm). In contrast to the result found at pH 4, NPs at pH 7 had a standard deviation of 0 nm, indicating greater stability and uniformity of particles. This particle size is appropriate, considering other standard studies, where the PLGA-NPs that were released directly into the mice synovial tissue had an average size of 265 nm. This size was suitable for the fast phagocytosis of NPs and easy penetration through the synovial membrane into the submembranous adipose tissue. Other studies have indicated that cells found in the synovial tissue are capable of phagocytosing latex particles with an average size of 240 nm. Since 255 nm falls between these standards, it can be considered an appropriate size for the NPs.

Likewise, zeta potential results were not directly related to changes in pH, as their values varied when the pH increased, indicating that pH did not have a direct influence on this measurement. Considering that a lower absolute value of zeta potential implies higher stability of NPs [cite], pH 4 would be the optimal pH based on zeta potential results. However, the high standard deviation (SD) at this pH (± 21.500 mV) might make it a less suitable candidate for RA treatment due to the poor consistency and charge instability of NPs. This could potentially lead to particle aggregation and inadequate functionalization. In fact, a dispersed distribution of zeta potential results was evident at pH 4, pH 5, and pH 8 (Figure. 3.11C, 3.11F, 3.11O) due to their high SD.

Although the absolute value of the zeta potential at pH 7 turned out to be one of the lowest, it had an SD of 0 mV, indicating a more uniform and stable charge distribution among particles at this point. In general, thanks to the low SD obtained at pH 7 for both size and zeta potential measurements, it is considered that the size and charge stability of PLGA-NPs would be feasible at a pH around 7. These results could be beneficial in treating moderate RA, as studies suggest that RA synovial fluid can have a pH ranging from 6.6 to 7.2. However, recent papers indicate that the pH in RA inflammatory tissue can decrease to as low as 6.0 or 6.5. For this reason, it is pertinent to conduct more detailed tests, varying the pH from 6 to 7, to assess how it affects the stability of NPs. This way, critical cases of Arthritis can be treated effectively.

From studying the physicochemical characteristics of the biomaterial, we can move on to study its biological properties. For this, the first thing that had to be done was to sterilize the biomaterial in order to obtain adequate results that would not be altered by microorganisms. From the sterility tests, due to the small significant difference between the absorbance of the samples analyzed, it can be concluded that the hydrogels were sterile, or at least mostly sterile, and therefore the sterilization process of the material was effective. The differences in the absorbances may be due to the composition of the material itself rather than the presence of microorganisms. In addition, this type of test is important because it allows testing for contamination of the material, which could lead to errors and false positives in subsequent biological tests [36]. The above applied in a laboratory setting, while in a medical setting sterilization is relevant to avoid infections and diseases that can harm the patient by the use of the biomaterial [36]. However, despite the effectiveness of the use of UV **r**ays for surface sterilization, in this case it is not the most app**r**opriate sterilization methodology because it only allows surface sterilization of the material and in this case not only the surface of the material will have contact with the human body, but also everything inside the hydrogel should also be sterilized as the MTX is released.

In the antibacterial activity of the material, it could be appreciated that the material did not present this type of activity, but, due to the verification of the sterility of the material, we cannot affirm that the material promotes the growth of bacteria. Although the material does not present its own antibacterial activity, this can be added by the use of antiseptics or antibiotics inside the hydrogel as ha**s** already been studied in different articles in this regard, such as the article by B. Pérez-Köhler et al [33], in which CHX was added to the hydrogel conferring antibacterial capacity fo**r** several days. Depending on the application the antibacterial activity can be more or less relevant, in the case of rheumatoid arthritis treatment an antibacterial activity could be effective to avoid possible infections in the application of the material due to the external environment during its use [35]. However, for the case of this application it is not required that the material has antibacterial properties to be functional, so it can be said that even if it does not have that quality, it is still appropriat**e**.

**Figure 4.1.**
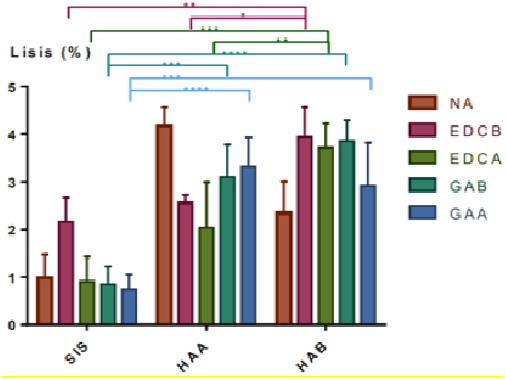
Percentage Lysis Red Blood Cells obtained from Oliveros Díaz [20].

Figure 4.1 shows the hemolysis results of a study based on the manufacture of regenerative dressings with the use of **H**A and SIS (porcine intestinal submucosa) hydrogels [20]. As is evident, all the samples containing HA had a percentage of lysis lower than 5%, which correlates with the results **ob**tained in Figure 3.9.

Regarding the hemolysis tests (Figure 3.9), it is evident that all samples had a percentage of RBC lysis lower than 5%, being the most hemolytic the GelHA (2% lysis). Therefore, it is considered that no hydrogel presented a hemolytic character (based on ISO 10 993-5 1992) [18]. This analysis is supported by other studies done with hyaluronic acid-based hydrogels [19,30], in which the hemolytic percentages were also below 5% [19] [31] (Figure 4.1). Thus, it can be stated that GelHA and GelHA-E hydrogels exhibit a low risk of red blood cell lysis. This is an essential point in this research, since one of the purposes of the drug used (MTX) is to reduce the degree of inflammation caused by rheumatoid arthritis, and therefore, an uncontrolled lysis of red blood cells near the synovial tissue could cause an undesired acute inflammation, which would be harmful to the patient, and consequently, this possible route of administration of the drug would have to be discarded.

**Figure 4.e.**
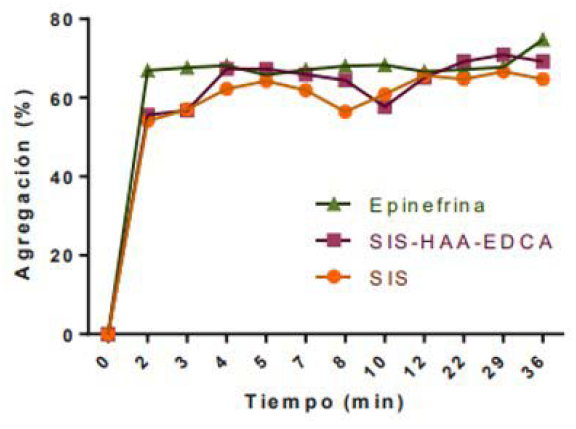
Platelet aggregation obtained from from Oliveros Díaz [20].

Now, as for the platelet aggregation test, the obtained aggregation percentages were 7% for GelHA and 6% for GelHA-E (Figure 3.9). According to these results, it is demonstrated that the GelHA and GelHA-E hydrogels do not have sufficiently high platelet aggregation percentages to form the platelet plug. However, according to the study performed by Oliveros Díaz, with a higher percentage of HA concentration a higher platelet aggregation of GelHA would be expected (Figure 4.2) [20]. Likewise, HA is hydrophilic. Therefore, when it reaches a point of equilibrium swelling, it begins to degrade, expelling the water molecules that it had captured [20]. This causes a greater release of the collagen that was going to react, and a greater aggregation of platelets [20]. Accordingly, it can be stated that the percentage of platelet aggregation in the gel is proportional to the HA concentration. Therefore, to increase the percentage of platelet aggregation, the concentration of HA in the hydrogel should be increased.

From the general procedure performed and presented, it can be observed that there were different limitations present both in the results obtained and in their analysis. One of the limitations, referring to the contact angle analysis, was the use of imprecise instruments in the image acquisition and the drop of the drop, which could affect the clarity of the angle analysis. Another limitation, regarding FTIR, was the resolution at which the data was taken, since there were points where it was not possible to appreciate differences with certain functional groups; this could also be due to the material itself, so it would not be possible to correct it. In the absorbance analysis it was not possible to perform the expected measurements, due to the rapid degradation of the hydrogel, so if a crosslinker with EDC were used, it would probably improve. Another limitation with respect to SEM is that liquid samples cannot be placed, which required the samples to go through a freeze-drying process as a preparation and there the morphology was altered so it was very difficult to know what it really was before this process. As for the sterilization process, the main limitation was the methodology used, which only allowed a superficial sterilization of the material. When analyzing the antibacterial activity, there were limitations in the effect caused by the material, so it would be necessary to replicate the procedure by adding antiseptics to the material. As for the platelet aggregation test, one limitation was the concentration of HA, since a smaller amount was used than was necessary to obtain a higher percentage of platelet aggregation. Likewise, another limitation was the reproducibility of the assay, since the results obtained could be considered as a trend, but for the results to be confirmed experimentally the methodology should be replicated at least 3 times on 3 different days. Finally, another experimental limitation of the platelet and hemolysis test was the limited resources available to repeat the assay or, more specifically, the blood available, since, when performing the first replication of the tests, all the available plasma was used.

Having said the above, one can move on to the bioethical analysis. As a bioethical principle of research, there is the responsibility as researchers to ensure the quality of the research, both in terms of interest, relevance and potential value, as well as in its ethical aspects [37]. A key bioethical consideration in this study is the in vitro characterization of hyaluronic acid- and eucalyptus-based hydrogels crosslinked with EDC. By thoroughly examining their physicochemical and biological properties, we aim to identify the optimal formulation prior to animal testing, thereby minimizing the use of animal models. This approach aligns with the 3Rs principle (Replacement, Reduction, and Refinement) in animal research. Furthermore, we propose the development of a subcutaneous model to evaluate the hydrogel’s performance when injected transdermally. This model could potentially be applied to assess systemic anti-inflammatory responses in a skin model or localized responses in a joint model incorporating synovial tissue. Such an approach would further reduce the need for extensive animal testing, contributing to more ethical and efficient research practices [38].

On the other hand, when analyzing the bioethical principles that should govern all research, it is possible to analyze that the principle of beneficence is fulfilled when considering that the initial research carried out has the principle of causing good to the greatest number of people currently suffering from rheumatoid arthritis. The principle of non-maleficence is fulfilled when considering that this study is in vitro and therefore the characterization of the hydrogel to know its potential for drug release is performed at this stage without the need to affect in any way any living being and optimizing the resource of products in the laboratory.

The principle of justice is fulfilled when considering that the main objective of the characterizations and evaluation of the biomaterial has its goal in a biomedical application that will benefit more than 0.24% of the world population [1] and more than 250,000 Colombians suffering from rheumatoid arthritis, 82% of whom are women [2]. If we start from the premise that we live in an unequal society, where patients affected by chronic, autoimmune, and systemic diseases, such as rheumatoid arthritis, are people who are in vulnerable conditions, this research and future work around it should be aimed at improving their quality of life.

In pre-clinical phases of the present research, the principle of autonomy will be applied, once the physicochemical characterization tests, as well as the in vitro and in vivo biological tests have been previously reported and repeated to guarantee the biosafety and technovigilance of the biomaterial [39]. When these tests are reported, the principle of autonomy can be guaranteed by formulating informed consents from individuals and patients who wish to be part of the clinical trial of this research. By means of the consents, it will be possible to inform about the procedures to be performed and the possible risks associated with the injection of a biologically evaluated material in vitro, and for the first time, in humans. The signature of the consents must always be done respecting the patient’s views and rights. In this way, the patient or individual volunteer can make an appropriate and conscientious decision [40].

## V. Conclusions and Future Work

In conclusion, this study generated a biomaterial based on hyaluronic acid and eucalyptus with MTX NPs as a possible treatment for rheumatoid arthritis. Thanks to spectroscopic, microscopic, and thermal characterization tests, the effectiveness of the crosslinking of the nanoparticles with GelHA-E was tested and it was possible to analyze aspects such as structure, hydrophilicity, solubility, resistance to degradation, and thermo-stability of hydrogels with and without eucalyptus subjected to different conditions. In addition, it was possible to test biological properties of the biomaterial such as antibacterial activity, hemolysis, and platelet aggregation.

The contact angle test revealed hydrophilic behavior in all samples, with average angles ranging between 15° and 21°. The addition of eucalyptus did not significantly alter the hydrogel’s hydrophilicity. Moreover, the HA gel demonstrated superior absorption in aqueous media, achieving 466.6% absorption. In contrast, MTX-PLGA-NPs-GelHA and GelHA-E showed the highest degradation rate, where MTX-PLGA-NPs-GelHA degraded almost completely. The above shows that, eucalyptus can enhance the shear thinning behavior of the hydrogel [29], however, the use of crosslinking agents such as EDC is considered necessary if it is desired to include eucalyptus in the material, with the purpose of increasing the degradation resistance in aqueous media and the swelling performance of the hydrogel in aqueous media.

Contrary to the absorption test, the TGA test indicated that, under high temperature conditions, the materials with eucalyptus have greater resistance to degradation. Indeed, the solutions without Gel-HA and eucalyptus (MTX-NPs-sol and NPs-sol) degraded rapidly near 300°C losing up to 3% of their weight, while the HA gels (GelHA-liq and GelHA-E-liq) were the most stable losing only 1% of their weight. This evidences the importance of the addition of Gel-HA in the stability of the biomaterial. Likewise, the compounds with eucalyptus (MTX-NPs-GELHA-E and GelHA-E-sol) had a more linear and controlled degradation than the other compounds, highlighting their usefulness to release the drug in a controlled manner when the material is subjected to high temperatures.

On the other hand, the FTIR test showed that GelHA-E-MTX-NP and GelHA-MTX-NP shared almost all their functional groups; however, an inverse relationship was evidenced between the proportions at which these groups were present in the eucalyptus gel and in the gel without eucalyptus. This, considering that the functional group with the highest absorbance in the HA-E-MTX-NP gel was the primary alcohol (absorbance of approximately 0.016), while in the HA-MTX-NP gel, the group with the highest absorbance was the double bond between carbons (absorbance of approximately 0.013). In general, both presented: OH groups present in PLGA and HA, methyl groups belonging to MTX and eucalyptus, ester groups belonging to PLGA and, finally, NH3 groups present in MTX. Thus, it is concluded that, the crosslinking of the hydrogel with the MTX nanoparticles was effective, since all the functional groups of the compounds were presented in the FTIR. In addition, a lamellar morphology with pores was identified when not using eucalyptus and without pores when using eucalyptus, as well as it was found that the biomaterials were not hemolytic (did not exceed 5%) nor was a high platelet aggregation obtained, which, in case of wanting to increase it, it will be necessary to use a higher concentration of the hydrogel.

pH changes did not significantly affect PLGA-NP size, as evidenced by zeta potential and size measurements. While the smallest diameter was observed at pH 4, it exhibited high variability. Conversely, pH 7 yielded the second smallest size with no standard deviation, indicating superior stability and uniformity. This size aligns with previous studies and is conducive to phagocytosis and tissue penetration. Zeta potential values fluctuated with pH; optimal values were observed at pH 4 but with high variability, whereas pH 7 demonstrated more consistent charge distribution. These findings suggest that a pH of approximately 7 may be optimal for maintaining size and charge stability of PLGA-NPs in moderate RA treatment.

Future investigations should focus on elucidating the relationship between eucalyptus concentration and hydrogel properties, specifically hydrophilicity, permeability, and crosslinking degree. Additionally, studies should aim to elucidate the mechanism by which eucalyptus influences the proportion of functional groups in the gels. To enhance the precision and reliability of results, it is imperative to employ more sophisticated analytical instruments, increase sample sizes, and perform additional replicates, thereby minimizing experimental error. Furthermore, comprehensive pH stability studies are essential to evaluate nanoparticle behavior across the range of pH levels characteristic of rheumatoid arthritis inflammatory tissue. These proposed studies will provide a more thorough understanding of the hydrogel system and its potential efficacy in RA treatment, ultimately contributing to the development of more effective therapeutic strategies.

## Suplemmentary Figures

**Figure a.3.1.**
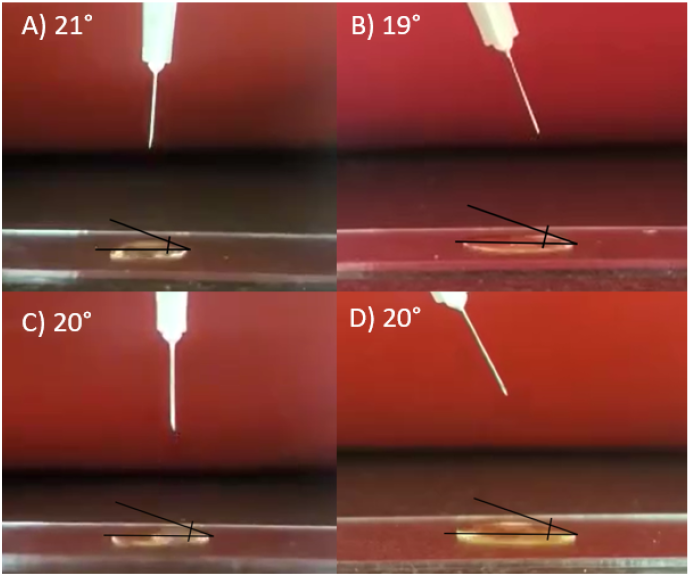
Contact angles obtained in the eucalyptus water hydrogel.

**Figure a.3.2.**
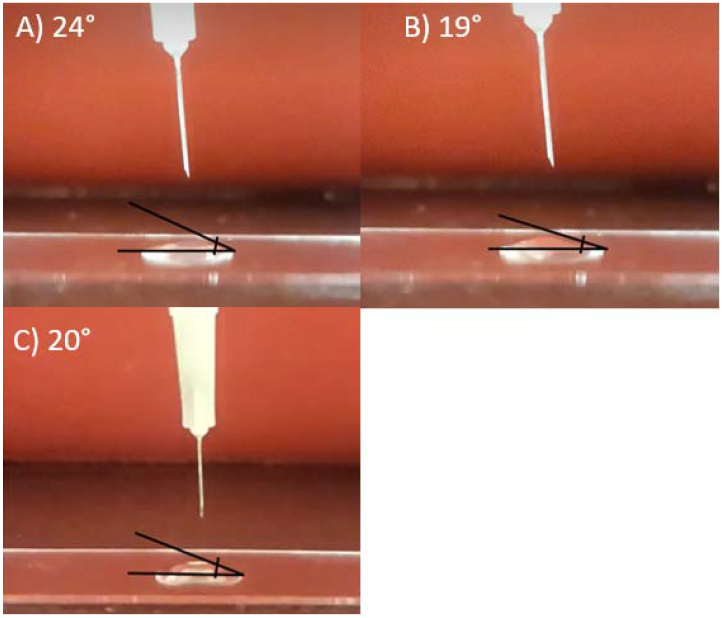
Contact angles obtained in GelHA hydrogels.

**Figure a.3.3.**
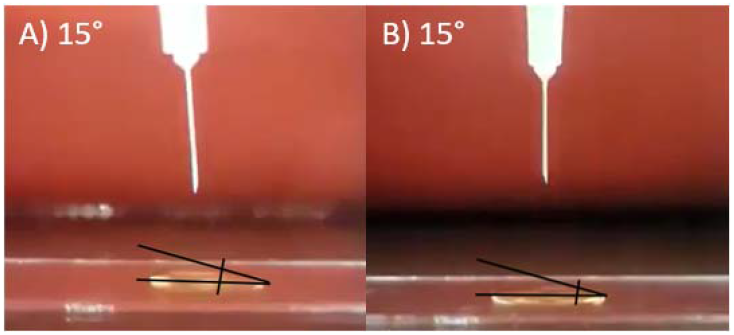
Contact angles obtained in GelHA-E hydrogels.

**Figure a.3.4.**
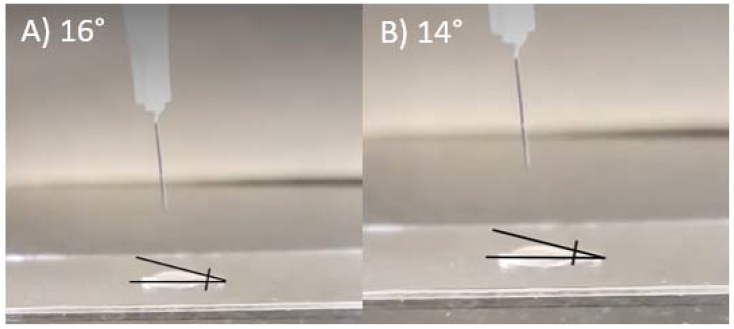
Contact angles obtained in the MTX-PLGA-NP-Gel-HA hydrogels.

**Figure a5.3.**
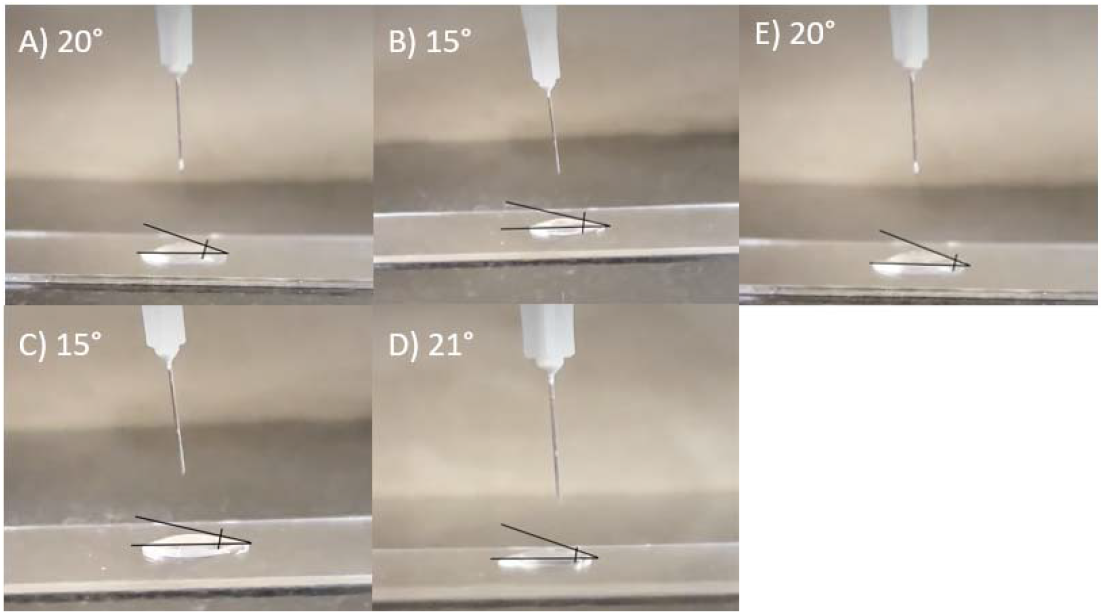
Contact angles obtained in the MTX loaded-PLGA NP and Gel-HA-E hydrogels Anexo A.4 Tabla de referencia para la preparación de extractos.

Anexo A.4 Tabla de referencia para la preparaciÓn de extractos

**Table A1.** Standard surface areas and volumes of extraction liquids.

| Thickness<br>mm | Extraction ratio<br>(surface area or mass/volume)<br>± 10 % | Forms of material |
| --- | --- | --- |
| < 0,5 | 6 cm<sup>2</sup>/ml | film, sheet, tubing wall |
| 0,5 to 1,0 | 3 cm<sup>2</sup>/ml | tubing wall, slab, small molded items |
| > 1,0 | 1,25 cm<sup>2</sup>/ml | larger molded item(s) |
| Irregularly shaped solid devices | 0,2 g/ml | powder, pellets, foam, non-absorbent, moulded items |
| Irregularly shaped porous devices (low-density materials) | 0,1 g/ml | membranes |
| NOTE While there are no standardized methods available at present for testing absorbents and hydrocolloids, the following is a suggested protocol:<br>Determine the "absorption capacity" of the material, i.e. the amount of extract liquid absorbed per gram of the material. The test sample shall be 0,1 g/ml beyond the absorptive capacity of the material. |  |  |

[19] ISO, E. 10993-12: 2004. Biological Evaluation of Medical Devices—Part, 12, 10993-10912 The table specifies the concentration of the extracts according to their shape and thickness.

